# Influence of Shea Tree (*Vitellaria paradoxa)* on Maize and Soybean Production

**DOI:** 10.1101/370155

**Authors:** Gertrude Ogwok, Peter O. Alele, Sarah Kizza

## Abstract

*Vitellaria paradoxa* provides many benefits to farmers within the Shea belt. However, increased threats to it necessitate its conservation, and one common approach is the practice of agroforestry. A number of studies have shown that Shea tree has influence on crop production, and yet, some of these studies were done using single season experiments or bioassays using mature Shea tree components. In this study, the seasonal influence of young and mature Shea trees on Maize and Soybean yields was investigated using field experiments in Otuke district of northern Uganda, where, Shea tree parklands are dominant and Maize and Soybean are used for food security and income. Our results show that there are differential responses of maize and soybean yield to rainy seasons and physiological differences of *Vitellaria paradoxa* treatment. We find yield reduction for maize more pronounced than yield reduction for soybeans under different Shea plants (Mature and Young) and for the two rainy seasons. We attribute the difference to the differential maize and soybean responses to *Vitellaria paradoxa* shading and its differential allelopathic inhibition of these crops. We recommend that Soybeans should be preferred to maize when planting under Shea canopy.

## 1. Introduction

The Shea tree (*Vitellaria paradoxa*) is an important parkland tree species indigenous to Africa, specifically occupying the Sudano-sahelian regions stretching from West Africa, across Central Africa to East Africa (Boffa, 2015). In Uganda, the Shea belt that occupies parts of Eastern and Northern Uganda predominantly has *Vitellaria paradoxa* sub species *nilotica* (Okullo, et al., 2012). The tree produces nuts that are processed to obtain Shea butter, which are of high economic value (Teklehaimanot, 2004). Shea butter has a wide range of uses including in the cosmetic and pharmaceutical industries (Gwali, Okullo, Eilu, Nakabonge, Nyeko, & Vuzi, 2012), and it contains important fatty acids including palmitic, stearic, oleic, linoleic and arachidic acids (Okullo J., et al., 2010).

Farmers in the Shea belt usually collect Shea nuts for domestic consumption and income and Shea butter is becoming an important foreign exchange commodity for countries where it is found (Rousseau, Gautier, & Wardell, 2015; Lovett & Haq, 2013). A number of scholars and stakeholders agree that there is need to conserve *Vitellaria paradoxa* due to its economic potential and threats to it (Buyinza & Okullo, 2015). These threats include burning the tree for charcoal, and large scale clearing of the Shea tree to pave way for mechanized agricultural production (Agea, Obua, Waiswa, Okia, & Okullo, 2010; Boffa, 2015; Buyinza & Okullo, 2015; Gwali, Okullo, Eilu, Nakabonge, Nyeko, & Vuzi, 2012). Also, increasing population pressure in the Shea belt means that there is no longer room for natural regeneration of the Shea tree (Okiror, Agea, Okia, & Okullo, 2012).

As a result, several conservation approaches are being encouraged by stakeholders including among others, intercropping Shea tree with annual agricultural crops. It is recommended that crops grown under Shea trees are shade tolerant (Boffa, 2015). More so, an increasing number of studies find that Shea tree has inhibitory effects on certain crops (Alamu & Aleem, 2014; Aleem, Alamu, & Olabode, 2014; Boffa, 2015; Folarin, Ogunkunle, Oyedeji, & Kolawole, 2015). The magnitude of the inhibition ranges from those that are not statistically significant and do not cause significant differences in yields, to those that are statistically significant leading to significant reduction in yields. The influence of planting season and crop productivity under Shea has not been widely reported. Majority of the studies on inhibitory effects of Shea tree are single season studies such as those of Aleem et al (2014) or bioassays that involve planting the experimental crops in soils incorporated with extracts of mature Shea tree components such as those of Folarin et al (2015). It is widely accepted that variations in seasons have significant influence on crop productivity (Boffa, 2015). Physiological stages of Shea is also expected to have significant influence on crop production. These influences are expected to vary with the crop in question, for instance, Alamu and Aleem, 2014; Aleem et al., 2014 reported differential responses of cowpea and maize to Shea tree. This study was therefore done to investigate the seasonal variations in the influence of *Vitellaria paradoxa* on production of *Zea mays* L. (Maize) and *Glycine max* (L.) *Merrill*. (Soybean) and investigate the influence of the physiological stage of the Shea on production of the two annual crops.

The two crops have been chosen for this study for two main reasons. First, both maize and soy beans are very important food security and income generating crops in the Uganda Shea belt. Secondly, these two crops were chosen for the study, due to differences in their physiology which is a big factor in their individual responses to the influence of *Vitellaria paradoxa*. Maize, for instance is a C4 plant, while soybean is a C3 plant. This means that the crops follow different photosynthetic pathways and are expected to exhibit different responses to the influence of *Vitellaria paradoxa* on their yields. We thus expect seasonal differences in the influence of *Vitellaria paradoxa* on both maize and soybeans, and a prominent influence of *Vitellaria paradoxa* on maize which, as a C4 plant is likely to be more responsive to variation in light intensity than soybean. We also expect different responses of both maize and soybean to different physiological stages of *Vitellaria paradoxa*, and that the influence of *Vitellaria paradoxa* on both maize and soybeans would be more pronounced for the mature Shea garden than young Shea garden.

## 2. Methods

### 2.1 Study Design and Data collection

#### Study Area

The study was conducted in the first and second rainy (planting) seasons in Opejal parish located in Okwang sub County, Otuke district, in Northern Uganda. It has a rainfall pattern with two rainy seasons from late March to May and July to November, with a long dry spell stretching from December to early March. The Average annual rainfall for the district varies between 1000 mm – 1600 mm. This rainfall is suitable for the production of both maize and soybeans. The mean temperature is between 22°C – 26°C. However, temperatures may be as high as 40°C during certain periods of the long dry season. Agriculture is the major source of livelihood in the district with major crops cultivated including; rice, groundnuts, sesame, soybeans, sorghum, beans, millets, pigeon peas and maize.

The natural vegetation is mainly savannah woodland with scattered trees dominated by Shea trees (*Vitellaria paradoxa).* Other prominent tree species includes *Terminalia, Cambretum spp*, *Ficus spp, Accacia spp* and *Phoenixma linareclinata.* Otuke district is generally flat or gently undulating with an altitude between 900 meters to 1500 meters above sea level, although much of the district lies above 1020 meters above sea level. The largest part of the district comprises of remnants of lowland surface, and is generally well drained except for some peripheral areas, that are occupied by poorly drained swamps. The district lies mostly in the Aswa water catchment that drains her wetlands in the south and west into river Aswa and Moroto respectively.

#### Experimental Design and data collection

The experiments involved three treatments of Mature Shea tree garden, Young Shea tree garden and a control garden that had no Shea tree. For purposes of this study, mature Shea trees were trees that were already producing nuts, while the young Shea trees had never produced nuts. The experiments involved planting both maize and soybeans in replicates of (1) four mature Shea gardens, (2) four young Shea gardens and (3) four Control gardens. Each treatment and control garden measured 10 × 15 meters. These were divided into sub-plots of 2.5 × 2.5 meters. In each treatment garden, a total of four sub-plots were planted with maize and another four sub-plots planted with soybeans in an alternating design (Figure 1). The planted plots were alternated with rest plots. This gives a total of 16 sub-plots for each treatment under maize and soybeans respectively. In season two, the alternating rest plots were planted alternately with maize and soybeans.

**Figure 1:**
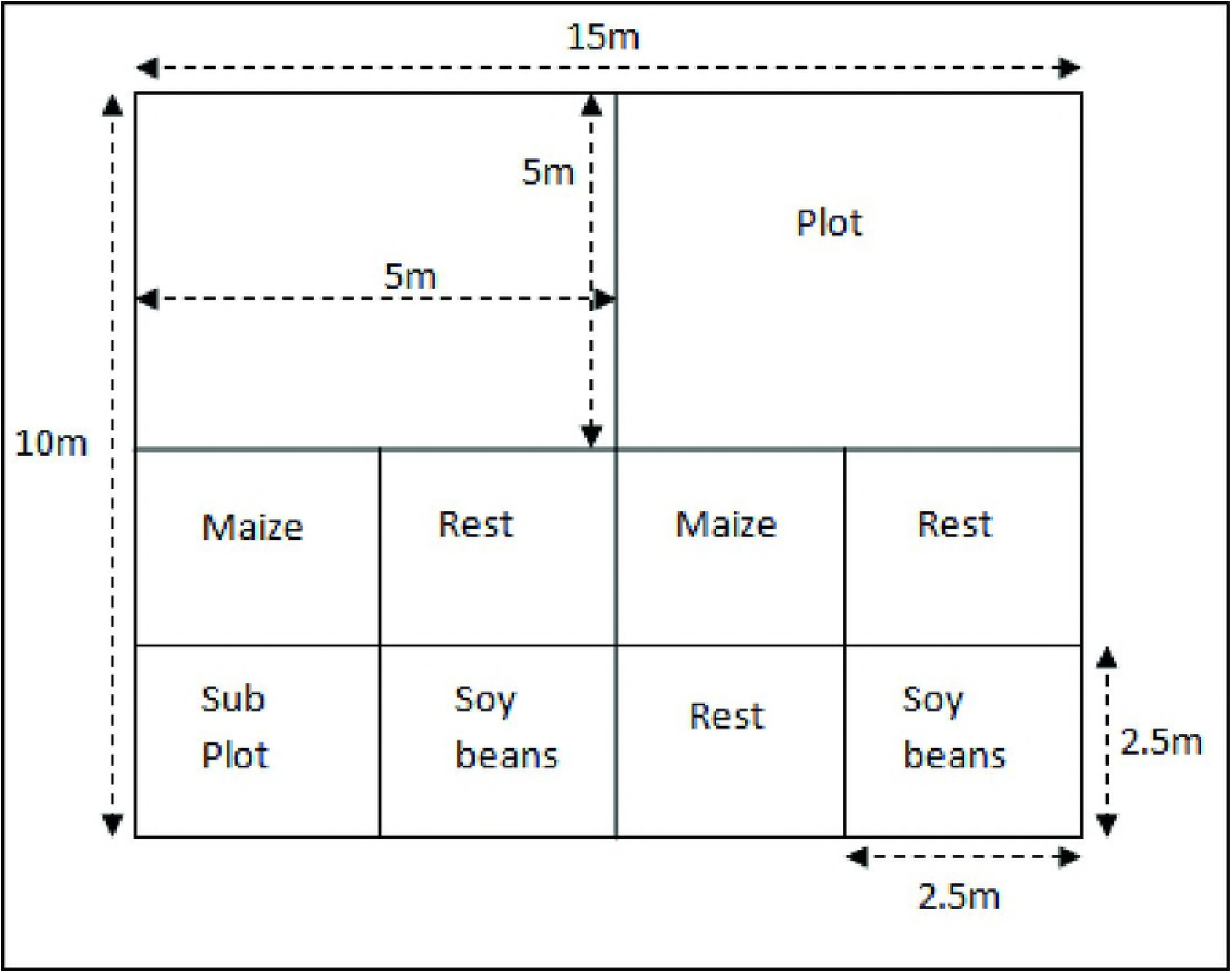
Treatment layout

The maize variety Longe 10H and soybean variety Maksoy 3N were used in this experiment. These varieties were chosen due to their drought resistance and high yielding potentials and are varieties that are recommended for the region. The agronomic practices undertaken for both maize and soybeans followed the standard practices that farmers in Otuke district follow. No inorganic fertilizers were used for all the two seasons. The crops were weeded twice as required. Maize and soybean yields after harvest, threshing and drying were weighed and recorded in grams for further analysis.

### 2.2 Data Analysis

Data on yields of maize and soybeans in both planting seasons were entered, cleaned and analyzed using statistical packages of Microsoft office excel 10 and SPSS version 20. The yield data was reported in kilograms per hectare. The data was subjected to Analysis of Variance (ANOVA) and post ANOVA to test for difference in yields within and between treatments at 5% level of significance. A yield decline index was also constructed to compare the yield difference between Maize and Soybean. The index was constructed by taking the yield from the control experiment as the base and comparing it with the yield from each of the two treatments as shown in the equation below.

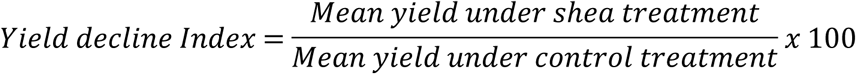

## 3. Results

### 3.1 Experimental Maize and Soybean yields

The results show variations in yield of maize and soybeans under the different treatments (Figure 2). Analysis of Variance found significant difference in mean yields for maize and soybean under the different treatment regimens for the two seasons (Table 1). Further analysis shows significant differences in mean yields for all the three treatments for maize, while there was no significant difference of soybean yield for mature and young Shea tree treatments. The mean soybean yields were however significantly different between the control and treatments and the results for season one was consistent with those of season two. However, there seems to be seasonal variations in yield within the treatments (Figure 1).

**Figure 2:**
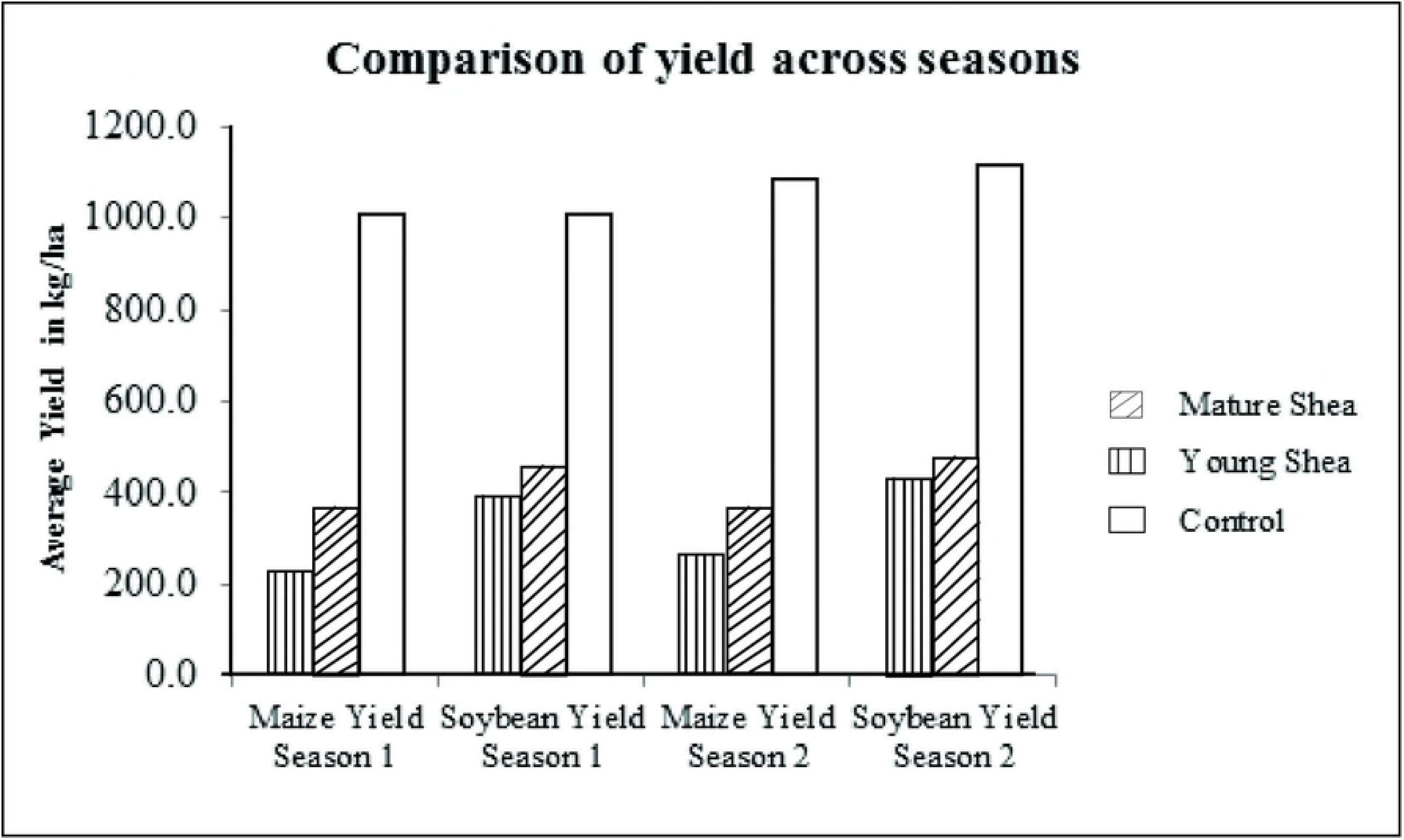
Maize and Soybean yields under different treatments across two seasons

**Table 1:** Variation in maize and soy bean yield among the treatments and control.

| Season One |  |  |  |  |  |
| --- | --- | --- | --- | --- | --- |
| Treatment | Number of Observations | Maize yield |  | Soy bean yield |  |
| | | Mean ± SD | p-value | Mean ± SD | p-value |
| Control garden | 16 | 633.06 ± 68.8<sup>c</sup> | 0.00 | 629.43 ± 159.70<sup>a</sup> | 0.00 |
| Young Shea | 16 | 231.62 ± 57.4<sup>b</sup> | 0.00 | 277.62 ± 57.60<sup>b</sup> | 0.00 |
| Mature Shea | 16 | 144.94 ± 23.8<sup>a</sup> | 0.00 | 170.25 ± 17.24<sup>b</sup> | 0.00 |
| Season Two |  |  |  |  |  |
|  | Number of Observations | Maize yield |  | Soybean yield |  |
| | | Mean ± SD | p-value | Mean ± SD | p-value |
| Control garden | 16 | 673.31 ± 51.06<sup>d</sup> | 0.00 | 700.18 ± 106.90<sup>d</sup> | 0.00 |
| Young Shea | 16 | 227.18 ± 48.24<sup>e</sup> | 0.00 | 297.25 ± 36.14<sup>e</sup> | 0.00 |
| Mature Shea | 16 | 170.25 ± 17.24<sup>f</sup> | 0.00 | 267.93 ± 17.17<sup>e</sup> | 0.00 |

*Note: SD Standard Deviation*
*Value in the same column with different superscripts are significantly different at $p < 0.05$*

### 3.2 Comparing maize and soybean yield response to treatment

Comparison of maize and soybean yields from the field experiments found that, overall, mature Shea tree treatment had the highest yield reduction for both maize and soybeans as compared to young Shea treatment. This reduction was more pronounced in the maize than in the soybeans, for instance, the average maize yield in season one for the mature Shea tree garden was only 23% of the average maize yield from the control garden while the average soybean yield in season one for the mature Shea tree garden was 42% of the average soybean yield from the control garden (Table 2). Similar results are seen for both maize and soybeans in season two.

**Table 2:** Comparison maize and soybean yield from the control with the treatments.

| Comparison | Yield decline Index (%) |  |  |  |
| --- | --- | --- | --- | --- |
|  | Season one |  | Season two |  |
|  | Maize | Soybean | Maize | Soybean |
| Mature Shea /Control Treatments | 23.1548 | 41.7235 | 25.4183 | 39.0404 |
| Young Shea/Control Treatments | 36.6053 | 46.3346 | 33.9325 | 43.1694 |

### 3.3 Comparing seasonal maize and soybean yield under treatment and control

Analysis of effect of season on the influence of *Vitellaria paradoxa* provides evidence of differential responses of maize and soybeans under the treatments. Specifically, maize yield under Mature Shea and Control gardens and soybean yield under mature Shea treatment were significantly different for the two planting seasons (Table 3).

**Table 3:**
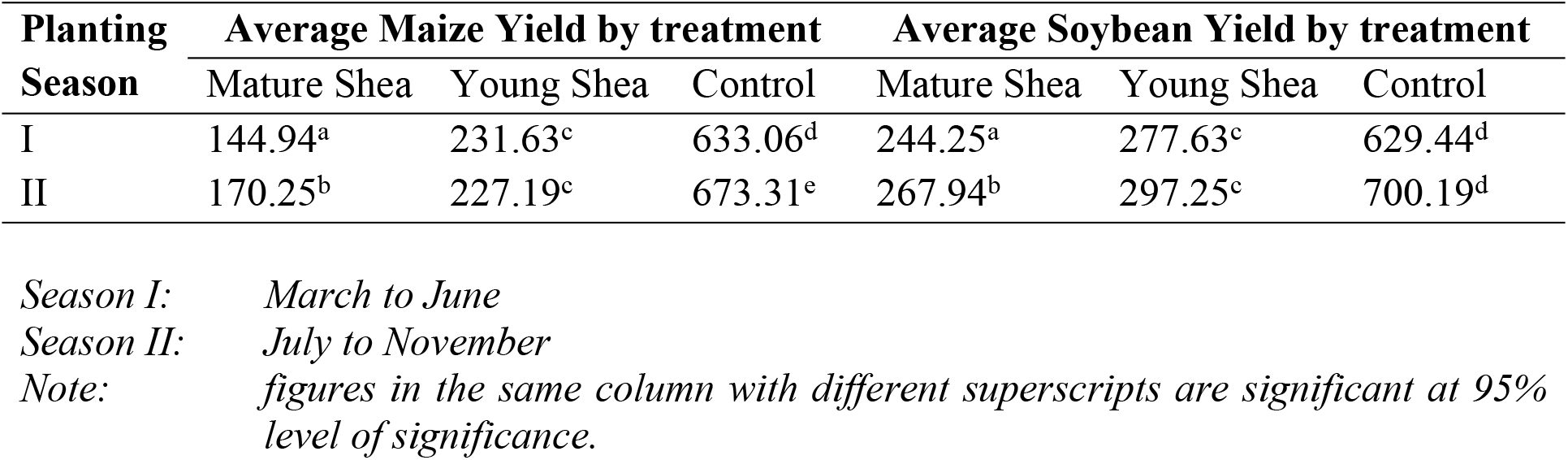
Seasonal Variation in Yield of Maize and Soybeans by Treatment.

## 4. Discussion

This study investigates the influence of seasonal and physiological stages of *Vitellaria paradoxa* on maize and soybean yields in Otuke district in northern Uganda. We find yield differences between the treatments, specifically; maize yield significantly different for all the treatments for the two seasons, while soybean yields not significantly different for mature Shea and young Shea treatments. We attribute this yield differences between the treatments to a number of factors; firstly, the response of maize and soybeans to different levels of shading as reported by (Suryanto, Putrab, Kurniawan, Suwignyo, & Sukirno, 2014). The difference between maize and soybean yield can also be attributed to the physiological differences between the two crops. For instance, Maize which is a C4 plant is very responsive to light intensity and temperature (Boffa, 1999). A C4 plant ‘s photosynthetic pathway uses different enzymes from those used by C3 plants. C4 plants are often called tropical or warm season plants. They reduce carbon dioxide captured photosynthesis to useable component by first converting carbon dioxide to oxaloacetate, which is a 4-carbon acid (Hopkins & Huner, 2009). Boffa (2015) indicated that C3 plants are less affected by Shea tree shading than C4 plants. This explains why our results show that the effect of *Vitellaria paradoxa* on Maize (a C4 plant) is more pronounced than in soybeans (a C3 plant).

We also attribute the yield decline due to *Vitellaria paradoxa*, to the presence of phytochemicals referred to as allelo chemical that have inhibitory effects on the growth and production of other plants. For instance, Aleem *et al*. (2014) and Alamu & Aleem (2014) in separate studies of cowpea and maize respectively reported that *Vitellaria paradoxa* had allelopathic effects on Cowpea and Maize. However, the influence of *Vitellaria paradoxa* on maize was more pronounced than in cowpea.

We also find that there were seasonal variations in the influence of *Vitellaria paradoxa* on maize and soybeans. Maize and soybeans yield in the second planting season were relatively higher than yields from the first season of planting. However, there was no significant influence of season on both maize and soybean yields under young Shea treatment and soybean under the control treatment. Mature Shea treatment showed significant (p<0.05) yield difference for both maize and soybean. This can be attributed to the seasonal variation in the chemical composition of *Vitellaria paradoxa.* Boffa (2015) reported that *Vitellaria paradoxa* exhibits phenology chronologically and geographically, while Byakagaba, Eilu, Okullo, & Mwavu, (2011) reported that Shea population structure and regeneration status depends partly on land management regimes. This means Shea tree chemical components that have inhibitory effects on growth and production of Maize and Soybeans vary across time and place. The differences in yields can therefore be attributed to differences in seasonal water availability.

### Limitations of the study

This study was carried out in two planting seasons. The results are therefore limited to situations similar to those of these two seasons. In case of very high weather variability to levels significantly different from these seasons, these results may not be applicable.

## 5. Conclusion

In this study, we investigated the influence of *Vitellaria paradoxa* on the yield of *Z. mays* and *G. Max.* Specifically, we expected to find varying influence of *Vitellaria paradoxa* on these two crops. We also expected to find differences in the seasonal influences of *Vitellaria paradoxa* on the two crops, and varying influence of mature Shea and young Shea on these two crops. It was expected that seasonal differences would lead to differences in crop responses under *Vitellaria paradoxa*, and that differences in the physiology of both maize and soybeans would lead to different responses of these crops to *Vitellaria paradoxa* under different physiological conditions and season. In line with our expectations, the study found that *V. paradoxa* influences maize and soybeans differently. Specifically, the influence was more pronounced in maize than in soybeans in both seasons. This can be attributed to the difference in response of maize and soybean to shading, and phytotoxic inhibition by other plants. The differential response of these crops can be attributed to differences in their physiology. We also find that mature Shea tree exhibited seasonal influence on both maize and soybean. This was attributed to the differences in the chemical composition of *V. paradoxa* especially with respect to allelo chemicals.

Our study provides a basis to recommend preferential planting of soy beans under Shea tree canopy as opposed to planting maize. This will help in furthering the effort to conserve the Shea tree (*V. paradoxa*), given its profound economic value

